# Computational design of toehold switches in eukaryotes and prokaryotes for efficient post-transcriptional control

**DOI:** 10.1101/2025.01.15.633215

**Authors:** Yehuda Landau, Matan Arbel, Daniel Benarroch, Jonathan Yoni Klein, Efi Moree, Tamir Tuller

## Abstract

The potential to design RNA molecules implementing translational regulated synthetic networks in eukaryotic cells can be harnessed to ramp up the development of various applications in biotechnology, synthetic biology, agriculture and personalized precision therapeutics. In prokaryotes, RNA based translation activation Toehold switches were demonstrated to achieve high fold change in throughput of a desired expression gene in response to different levels of both exogenous and endogenous target trigger RNA molecules. However, low fold changes were reported for such switches applying similar design architecture for eukaryotes. We developed Triggate, a computational pipeline which implements different algorithms for the efficient design and generation of higher on/off ratio optimized toehold switches in prokaryotes and eukaryotes. We also present here a new design architecture for eukaryotic toehold switches which is leaner and simpler. Using our tool and new architecture we report results with a much higher on/off ratio (up to around 13) than achieved before for mRNA toehold switches for eukaryotes.

Triggate is available as a web application at http://www.cs.tau.ac.il/~tamirtul/Triggate.

## Introduction

Riboswitches are regulatory gene expression elements, designed primarily to modulate transcription termination or translation initiation of mRNAs. One advantage of riboswitches is their relatively simple design being composed of two domains, a ligand-binding aptamer domain, and an expression platform. The aptamer domain typically resides in the 5’ untranslated region (UTR) and upon specific ligand binding dictates conformational change and regulation of expression [1], [2]. The standard riboswitches commonly found in prokaryotes modulate translation efficiency by switching between two bi-stable structural states: one that sequesters the ribosomal binding site (RBS), typically in a basic hairpin secondary structure, resulting in lower efficiency of ribosomal recruitment, and the other in which the RBS is exposed and ribosomal recruitment is more efficient [3].

Following the discovery of riboswitches in nature, synthetic biology researchers were inspired to mimic that behavior [4], [5].In nature, the nearly 40 different classes of riboswitches discovered so far, are characterized by aptamer domain folding into ligand-binding pockets for specific small molecule ligands [2].

However, relying on RNA-RNA interactions to modulate aptamer folding allows the design of sequence-specific RNA molecular switches. Under the reasonable assumptions, that the secondary structure already determines a stable three-dimensional architecture and is predictable according to Watson-Crick base pairing [6], computational models and algorithms based on thermodynamic equilibrium free energy experiments and dynamic programming optimizations were developed as a set of tools for RNA structural and RNA-RNA interaction modeling [7].

Engineering mRNA switches designed to be triggered by other RNA molecules can be greatly beneficial for synthetic biology applications. Since different cell types, cell states, and organisms have typical transcriptomic profiles [8], [9], [10], the ability to design switches triggered by the presence of specific mRNA markers allows engineering specific targeting methods for therapeutic and diagnostic purposes.

The first-generation synthetic switches for the regulation of translation initiation involving RNA-RNA interactions were designed for prokaryotes, mimicking the natural general architecture [4]. The best demonstrated dynamic range was up to 50-fold, which was one order of magnitude lower than demonstrated for protein-based transcriptional regulators.

A decade later a new generation of synthetic RNA-based switches for prokaryotes were presented using a de-novo architectural approach, called toehold switches. The major design changes included adding a linear sequence (an unstructured RNA segment is termed linear) at the beginning of the 5’ UTR, before the hairpin structure, intended for trigger binding with a linear trigger resulting in more thermodynamically favored linear-linear interaction, sequestering the RBS sequence and the start codon in a stem-loop. These switches achieved a dynamic range of over 400 [11]. Based on these results the applicability of designing toehold-based biosensors was demonstrated [12].

In contrast to the high dynamic range demonstrated for toehold-based switches for prokaryotes, the transition of the key architectural design pillar of RBS sequestration to Kozak sequestration for eukaryotes achieved a dynamic range of only up to 2-fold [13]. The ability to use toehold-based switches for sensing miRNA as triggers has also been demonstrated. miRNAs were shown to be involved in a range of biological processes, including development, cell proliferation and differentiation, as well as being involved in human diseases including cancer [14], [15].The low dynamic range achieved for Kozak and start codon (AUG) sequestration in eukaryotes can be attributed to the different mechanism of translation initiation, which canonically starts with the small ribosomal subunit scanning the mRNA from the 5’ cap rather than being directly recruited to the Kozak sequence and the following start codon [16]. In pursuit of enhancing the induction of a eukaryotic translational switch by bypassing the 5′ cap scanning process, a switch relying on noncanonical translation initiation was developed by modifying a known Internal ribosome entry site architecture, achieving up to 16-fold activation for a synthetic RNA trigger and up to about 5-fold for endogenous transcripts [17].There are additional approaches for regulating translation in a condition specific manner (e.g. based on ribozyme [18] or RNA binding proteins [19], [20] and RNA binding proteins combined with miRNAs [21]) but each of these approaches has its specific limitations and disadvantages.

Existing software tools for the design of RNA-based toehold switches allow the assessment of switch and trigger candidates using energy calculations and machine learning models but are based on limited data that has mostly been tested in prokaryotes and are biased toward specific architectures [22], [23], [24]

In this study, we developed a computational pipeline and tool for designing toehold switches in prokaryotes and eukaryotes, including novel architectures that better fit eukaryotes. We demonstrated for the first time that we can achieve dynamic range of around 13-fold for eukaryotes when using mRNA as a trigger.

## Results

### Pipeline for computational design of toehold

We have developed a pipeline for the computational design of toeholds.

#### Cross domains architectures

Our tool implements cross domain toehold design including different architectures for prokaryotes and eukaryotes. We implemented one general architecture for prokaryotes which is the “canonical” one used by previous works [11] (Fig. 1A), a similar architecture adapted for eukaryotes (Fig. 1B), and a de-novo architecture for eukaryotes (Fig. 1C).

**Fig. 1:**
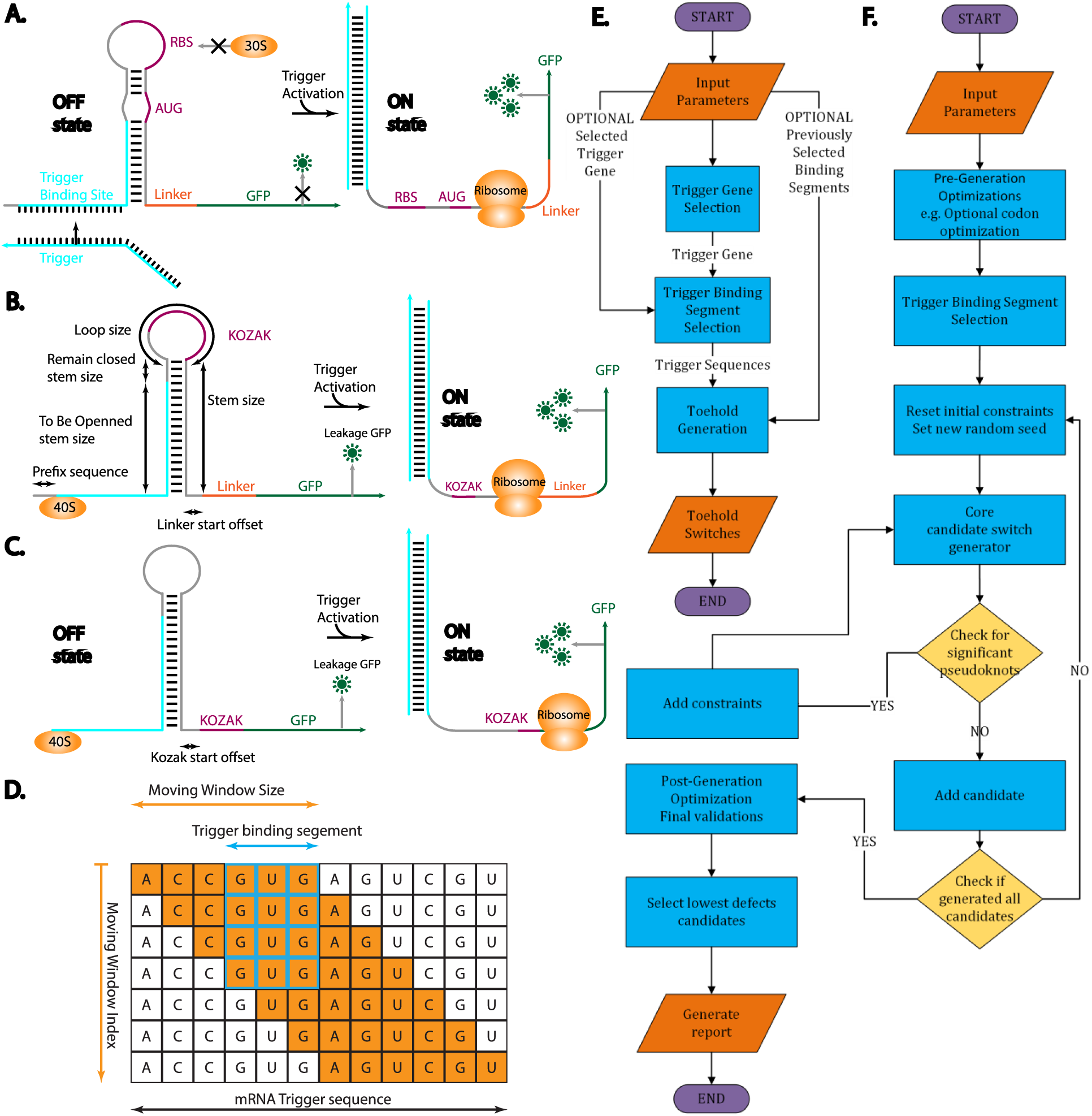
Toehold switch architectures activation diagrams, flow charts and algorithms of the computational tool. **A.** Ordinary prokaryotes toehold switch design and activation diagram. **B.** Ordinary eukaryote toehold switch design and activation diagram. **C.** De-novo eukaryotes toehold switch design (placing Kozak after stem) and activation diagram. **D.** Algorithm for segment accessibility approximation in the trigger selection module using a moving window algorithm. **E.** High-level flow chart describing the computational pipeline. **F.** Flow chart describing the toehold generation module of the pipeline.

#### mRNA as a trigger

Our tool allows the usage of a mRNA as a trigger, by implementing algorithms (Fig. 1D) to find the most probable unfolded segment of a trigger mRNA to serve as an optimal candidate to bind with the toehold switch binding site and change its state from off to on.

#### Outline of the pipeline

The toehold generation process can be divided into 3 main stages, implemented by running through a sequential combination of processing sub modules, each using the previous output as its input (Fig. 1E,F). The first stage is the selection of a trigger mRNA gene. In the current version of our tool, we assume the user has already selected a trigger mRNA molecule. The second stage is finding sub-segments in the selected trigger gene to serve as an activation sequence for the toehold switch. This stage may be optionally skipped when a user specifies a previously chosen trigger segment. The third and final stage is the generation of a toehold sequence according to specified parameters and requirements optimized to be activated by the trigger sequence selected in the previous stage or provided directly by the user.

#### Special design features and optimizations

Previous work has focused on optimization of the on/off ratio as well as preferring lower leakage designs (low off signal). However, for the potential implementation of the technology both for diagnostic and therapeutic applications the on-state signal carrying on the desired effect of the switch-regulated gene is of crucial importance. To optimize the expression level of the gene we have implemented translation optimization sub-modules, mainly implementing codon bias for both the expressed gene and linker when used. Gene codon optimization is done by exploiting codon bias adaptation toward codons that statistically appear more frequently in highly expressed genes [25] of the selected organism. The codons were also chosen while considering the optimality of the designed switch.

#### Cross organism and cell type parameters

The mentioned translation optimization is dependent on statistical data collected for a specific organism type. We have implemented a special cross organism optimization database for a few model organisms and cell-lines including *E. coli*, *S. cerevisiae,* HEK293 cell line and more. Our database also holds different optimized RBS and Kozak sequences appropriate for those model organisms, but to allow the selection of other organisms, our tool implements the option for customized usage by directly setting these parameters.

#### Measurement parameters

Our tool sets a “canonical” set of default measurements that have been successfully used in previous works, while allowing the advanced user to diverge from these default values of the stem size, loop size, binding site size, as well as many other parameters, encouraging the simple exploration of different measurements within the few prescribed architectures.

### Heterologous mRNA targeting

Using our computational pipeline, we generated 8 toehold switch variants to be tested in yeast. The 8 variants are divided into two groups, each focusing on one of the two major architectural designs for eukaryotes available in our tool: 4 variants using our novel architecture which places the Kozak sequence after the hairpin stem, and 4 variants implementing the architecture with the Kozak placed inside the hairpin loop, mimicking the architecture commonly used for prokaryotes to sequester the RBS [11].

The expression gene tested was GFP and the trigger gene selected was mCherry. We tested two versions of the mCherry gene as targets, with the same amino acid sequence but different codon adaptation index scores. One version of mCherry is the same sequence previously used by some other works [11], [12] originally tested in prokaryotes, and the other version is codon translation optimized for yeast. For the prokaryotic version of mCherry we have used the same window segment for trigger binding used successfully by a previous work, and for the yeast optimized version we have used our algorithm to calculate the optimal estimated segment for trigger binding. Each of the two architecture groups tested were thus further divided into 2 sub-groups, each tested against one of the two mCherry mRNA trigger versions.

We then generated several candidates for each of the 4 sub-groups and selected two of them with the highest predicted metric score according to our computational model.

All variants were generated with similar measurements to those previously demonstrated to achieve optimal on/off ratio for prokaryotes in related works [11]. We have used codon bias relaxation method of the first 6 gene expression codons for enabling more degrees of freedom toward directing selection of candidate with higher predicted metric scores.

To validate our algorithm, we constructed a simple and straight forward experiment to exhibit the algorithm capabilities to construct de-novo toeholds on demand. In our system, one plasmid expresses the mCherry ‘trigger’ RNA and a second plasmid expresses GFP coupled to different toeholds. An illustration of the experiment is shown in Fig. 2A. We constructed a set of 8 such plasmids with different toeholds in yeast and then transformed them with the ‘trigger’ plasmid, allowing us to examine GFP fluorescence levels with and without the trigger on the same background and extrapolate the ON/OFF ratio of the toehold. This system also allowed us to validate the presence of the trigger by fluorescence. In addition to quantifying the ON/OFF ratio, we examined how the presence of the toehold affects the absolute expression level of the coupled gene, by comparing the GFP fluorescence levels of the toehold constructs to an ‘open GFP’ construct which lacks a toehold. While some of the toeholds demonstrated high activation in the OFF state and some demonstrated low ON levels, 4 of the 8 toeholds performed well, shutting off expression when not in the presence of the trigger but allowing expression close to that of an ‘open GFP’ in the presence of the trigger (Fig. 2B). Some of the toeholds exhibited ON/OFF ratios over 10, and as high as 13 (Fig. 2C). A possible partial explanation for the fact that none of the toeholds reached an ‘open GFP’ fluorescence level is the fact that the 5’ UTR region added to the mRNAs to include the toehold has pushed the gene further away from the promotor, which may lower its overall expression levels.

**Fig. 2:**
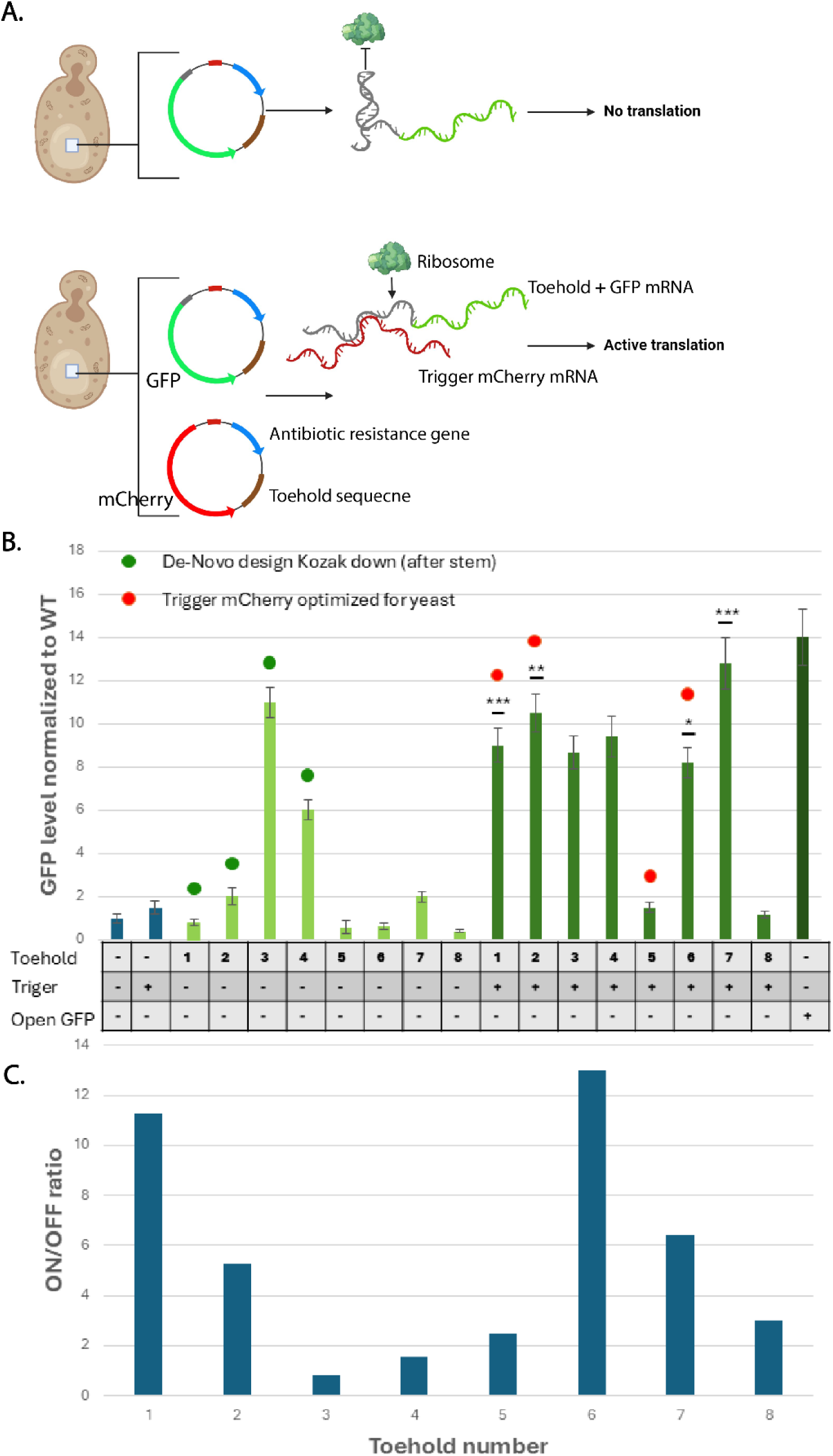
Toehold switches with heterologous mRNA triggers experiment design and activity results. **A.** An illustration of yeast cells with either toehold alone or toehold with trigger. **B.** GFP fluorescence levels (normalized to WT-Yeast without any plasmid) of yeast cells with either toehold alone or with trigger. The data shown is an average of three repeats, error bars showing standard deviation from the average. P-values signified by *, **, *** represent the ON state being significantly higher than the OFF state - *, P <0.01, **, P <0.0075, ***, P <0.005. **C**. ON/OFF ratio of the different toeholds. While some demonstrated little sensitivity to the presence of the trigger, almost 50% of the toeholds worked and some worked with more than 10-fold induction level.

### Endogenous mRNA targeting

The previous experiment demonstrated the ability of the toeholds to selectively express GFP in the presence of a trigger introduced on a plasmid. Next, we wanted to demonstrate the ability to target endogenous genes in eukaryotes as this would better illustrate the usage of regulated translation by endogenous mRNA in different biotechnological applications. To achieve this, we designed toehold switches to target heat-shock protein mRNAs in yeast, which are upregulated only when yeast cells are exposed to elevated temperatures (Fig. 3A). HSP proteins were selected as a target for this experiment because they are relatively simple to induce, well researched, and have a high difference in expression between the states. In this setup, under normal conditions, where the target mRNA is not transcribed, the switches remain in their locked OFF configuration. In contrast, when yeast cells undergo heat stress, the target mRNA is transcribed, prompting our toehold switches to transition from their locked OFF configuration to an unlocked ON state.

**Fig. 3:**
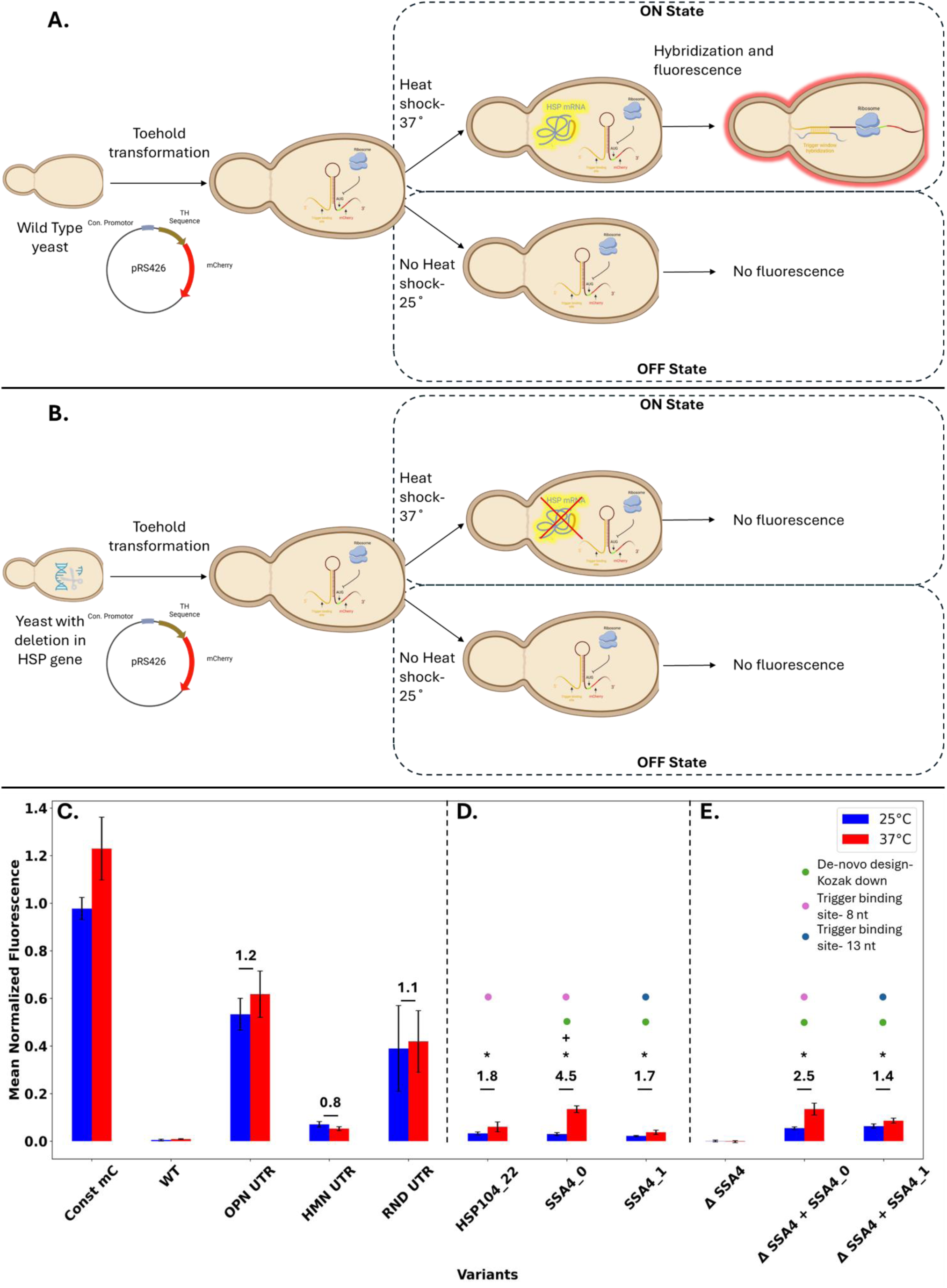
Toehold switches with endogenous mRNA triggers experiment design and activity results. **A.** Experiment setup targeting endogenous heat shock genes. **B.** Experiment setup with HSP gene deletion. **For C, D, E -** mCherry signal intensity (excitation: 580 nm, emission: 620 nm) was normalized to cell density (OD600). The mean normalized fluorescence was calculated by dividing all measurements within a plate to the measurement for constitutive mCherry at 25°C and then subtracting the mean signal for the corresponding temperature in the WT strain. Error bars represent one standard deviation in each direction. ON/OFF ratio of selected toeholds is shown above the respective bars. Variant names are XXX_YY (X=target gene name, Y-variant id). P-values signified by * represent the ON state being significantly higher than the OFF state - *, P <0.01. P-values signified by + represent the ON/OFF ratio of the variant being significantly higher than the ON/OFF ratio of the UTR controls (Human, Open, Random) and constitutive mCherry: +,P <0.01. **C**: Control variants. **D**: Tested Toehold variants. **E**: Deletion strain results. Δ SSA4-yeast strain with a deletion in the SSA4 gene-no Toehold. Δ SSA4 + SSA4_X are yeast variant with deletion in SSA4 gene and with inserted toehold variant.

We selected 2 heat shock proteins (HSP104, SSA4) that have been demonstrated to have high expression levels during heat stress, and low expression under normal conditions [26], [27], [28], [29].

For each of the 2 targets, we evaluated four distinct toehold topologies, which represent combinations of two toehold components: the Kozak location, as mentioned before, and the number of nucleotides in the trigger binding site that are designed to be unhybridized in the locked state (8/13 out of 25 nucleotides). We selected 3 of the variants to examine experimentally. In this experiment, the toehold was coupled to the mCherry gene, and mCherry fluorescence levels were quantified with and without induction of the heat shock proteins (Fig. 3A,D).

While some residual leakiness was observed in the non-induced state compared to the wild-type strain, the results demonstrate that some of our toehold switches effectively respond to their target during heat stress. The SSA4_0 variant specifically stands out, exhibiting a substantial ON/OFF ratio exceeding 4.

To control the changes in the 5’ UTR of the expressed gene, mCherry fluorescence was also examined with constructs that contain a randomized UTR sequence, a human-derived UTR sequence, and an ‘Open UTR’ instead of the toehold. These constructs are not expected to regulate mCherry expression. While all the tested toehold switches demonstrate a higher ON/OFF ratio, the controls also show some difference in expression between the states (Fig. 3C, Fig. Supplementary figure 1). We hypothesize that this might be due to the change in temperature affecting the mRNA expression and folding directly. This phenomenon might affect the activation levels of the toeholds as well, meaning some of the difference in signal between the induced and uninduced conditions might be due to the change in temperature rather than opening of the toehold due to trigger binding. To control for this possibility, we examined mCherry fluorescence levels for the SSA4-targeted toeholds in strains lacking the SSA4 gene. Since the trigger is absent, the ON/OFF ratio in these strains corresponds solely to temperature-related effects (Fig. 3B,E).

The results for the strains with SSA4 deletion and toeholds suggest that exposure to heat appears to have some effect on the increase in normalized mCherry signal intensity, even in the absence of the trigger. However, when comparing the ON/OFF ratio between variants with and without SSA4 deletion, the ON/OFF ratio in variants with the trigger was significantly higher than that observed in the variants without triggers. This indicates that the observed ON/OFF ratio largely reflects the functionality of the toehold, successfully transitioning from the OFF to ON state in response to the trigger, rather than being solely due to a heat-induced effect.

### A user-friendly web tool

Our web tool user interface is implemented in a single form (Fig. 4A,B). We released a user-friendly web tool to enable the usage of our pipeline. A single dynamic form enables the selection of different domains and architectures, different organisms for specific optimizations, the providing of trigger mRNA and expression gene, and the setting of many customizable design measurements. Finally, a file with an assortment of model optimal candidates is returned as well as a visual report. More details appear in the methods section and in the tool website.

**Fig. 4:**
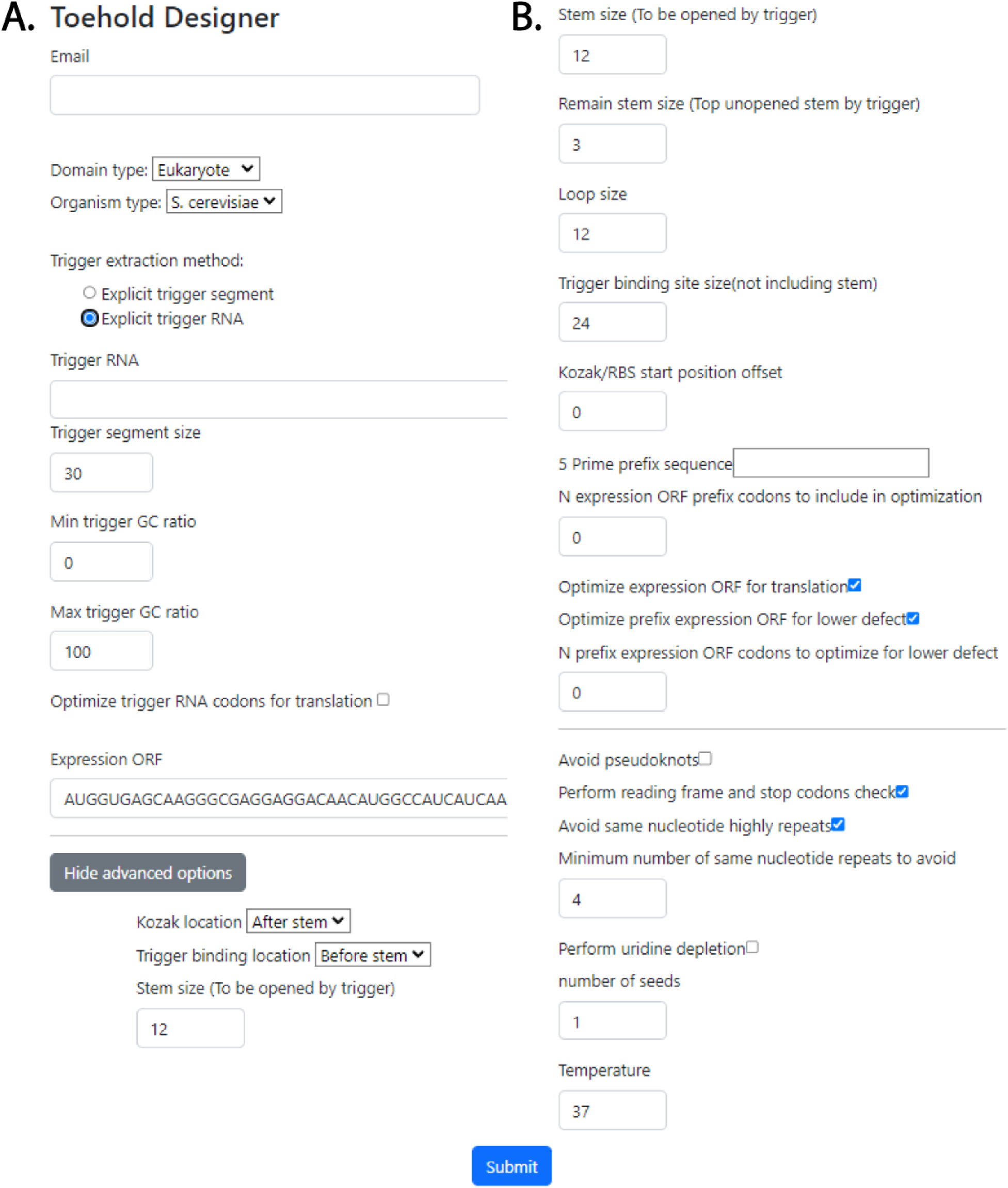
Web-based user interface for the computational pipeline. **A.** Example of an upper form screenshot. **B.** Example of a lower form screenshot.

## Discussion

Since the ON/OFF ratios for toehold switches tested in prokaryotes were much higher than those previously demonstrated in eukaryotes, we hypothesized that though some of the fundamental molecular processes are common to both domains (mostly those concerning the toehold branch migration process), many translation-related processes act differently in eukaryotes.

One of the major translation-related differences is the translation initiation process, which is manifested by the difference in functionality of the small ribosomal subunit.

In prokaryotes, the small subunit binds directly to the RBS sequence of an mRNA, signaling the beginning of translation initiation. Thus, the RBS serves as an ideal candidate for translation down-regulation through its sequestration when localized in the toehold switch stem-loop (Fig. 1A).

In eukaryotes, however, the canonical (without IRES) initiation process is characterized by the small ribosomal subunit scanning of the mRNA from the beginning of the 5’ UTR, starting in the 5’ cap. Many eukaryotes initiation factors (eIF) are involved in this process, some of which (e.g. eIF4A) are responsible for helicase unzipping of the double-stranded segments [16] Since translation initiation does not depend on direct recruitment of the small subunit to the Kozak sequence [16], [30], [31], [32], sequestering the Kozak sequence in a stem-loop (Fig. 1B) may not efficiently prevent translation. Moreover, the high efficiency of the initiation complex, and specifically the helicase eIF4A which is stimulated by other eIFs (eIF4G, eIF4B), in unwinding secondary structures during mRNA recruitment and scanning [33], [34] can explain the high off state translation rate in eukaryotes, leading to lower on/off ratios.

One of the novelties of our work is the presentation of new toehold switch design architectures demonstrating a change in the placement of the Kozak sequence. In contrast to previously demonstrated research and toehold library design in both prokaryotes [11], [12] and eukaryotes [13] we have decided, following our mentioned hypothesis, to construct a novel architecture which places the Kozak sequence after the designed hairpin structure followed by the expression gene (Fig. 1C). In such a design strategy, we also avoid the need for adding a linker, which was often necessary in previous designs since translation would begin within the stem of the hairpin, which must resemble a segment of the trigger sequence. Another benefit of this architecture is the relaxation of the constraint to avoid an in frame stop codon in the stem following the Kozak sequence, which has posed a constraint on the selected trigger segment.

The novel positioning of the Kozak sequence after the hairpin’s stem has introduced a new flexibility in the design architecture that could be exploited intelligently, leading to another novel feature. Placing the Kozak sequence at some distance after the stem (Fig. 1C) adds a degree of freedom to the algorithm, allowing the optimization of the pre-Kozak sequence in such a way that decreases the local binding affinity of the trigger binding site (expected optimally to be a single-stranded segment ready for the trigger) to the Kozak sequence and the downstream coding sequence (Fig. 5A).

**Fig. 5:**
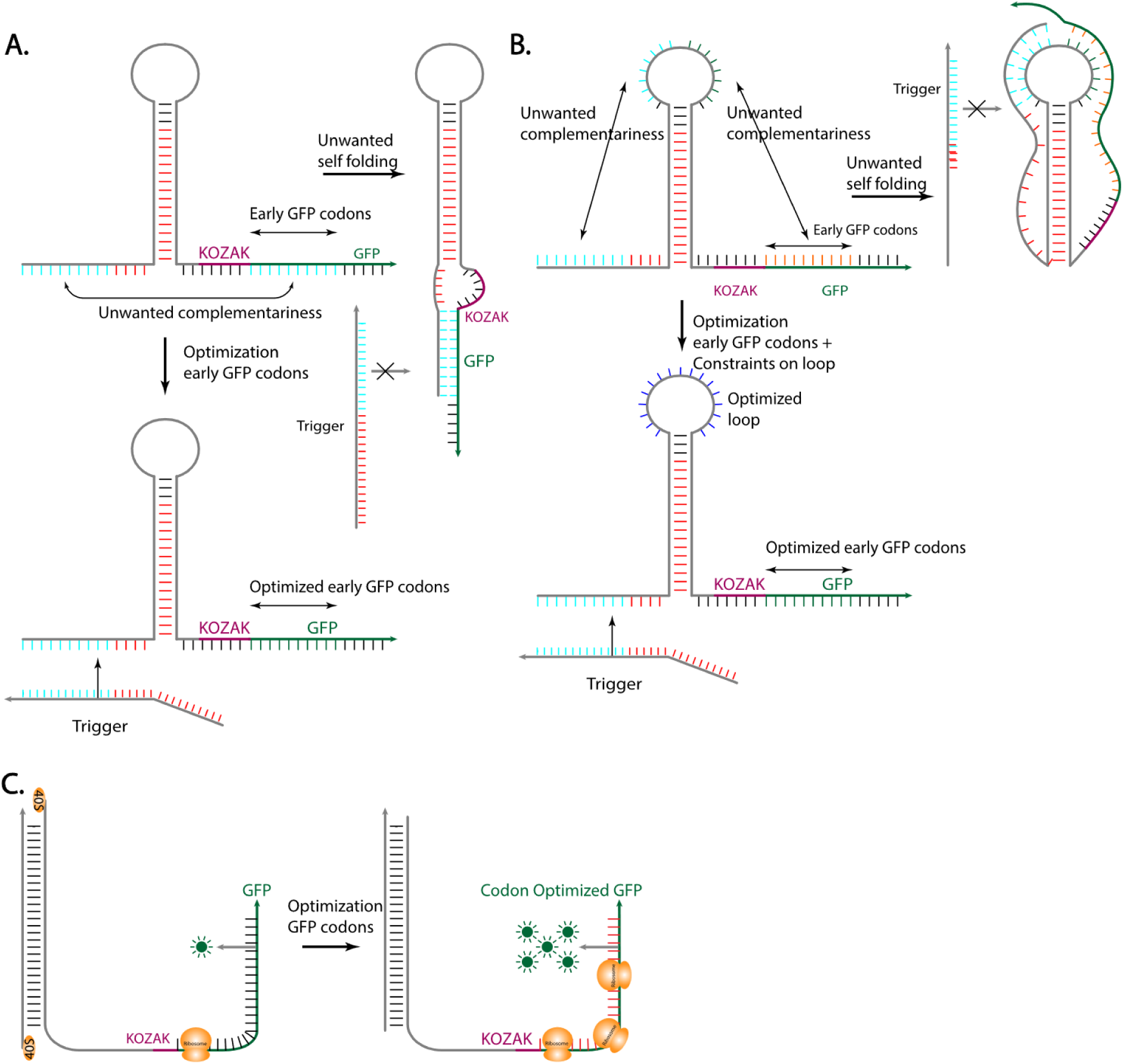
Toehold switch algorithms illustrations. **A.** Early codons optimization for lower ensemble defect score avoiding unwanted linear-linear interactions. **B.** Pseudoknots avoidance algorithm for loop-linear interactions removal. **C.** All ORF codons optimization for higher translation throughput.

In the first experiments, we generated 8 variants of toehold switches for eukaryotes using our computational tool and tested them in yeast. For 4 of them we used a yeast optimized version of mCherry, for which we selected a trigger segment using our algorithm, thus implicitly validating its integrity. Nevertheless, future studies on large libraries of toehold variants will be required to fully validate the algorithm. Examining our de-novo architecture placing the Kozak after the stem and not requiring a linker, showed similar on/off ratio to those with Kozak placement in the loop. Though we did not get superior results with our de-novo architecture, the fact it achieved similar results validates our hypothesis that the sequestration method has no positive effect. As a result, we recommend using our de-novo architecture, which achieves similar efficiency while not requiring a linker which is very important for many biotechnological applications; in addition, the new architecture releases some algorithmic constraints as explained above.

The much higher ON/OFF ratio we achieved (around 13-fold) compared to prior tested canonical mRNA designed toehold switches for eukaryotes (around 2-fold) can be attributed to many features we have used. We used codon optimization to improve the translation of the translated mRNA and codon bias optimization of the early expression gene codons for achieving lower defect score as well as pseudoknot avoidance. The relative contribution of each feature could only be determined in future research tested on larger libraries.

The second experiment marks a significant achievement in demonstrating the feasibility of using toehold switches to target endogenous genes in eukaryotes. By successfully designing toehold switches that respond specifically to native heat-shock protein (HSP) mRNAs, we have shown that it is possible to achieve selective gene expression without relying on exogenous triggers. triggers.While the ON/Off ratios achieved in this experiment are lower than the values shown in Fig. 2C, it is to be expected since the trigger in this experiment is an endogenous gene that is still somewhat expressed in the OFF state and not as highly expressed in the ON state as an artificially inserted gene on a high copy-number plasmid.

This capability is particularly novel, as it expands the applicability of toehold technology beyond prokaryotes and artificially introduced sequences, opening new possibilities for targeted gene regulation in more complex organisms. The successful ON/OFF response observed in variants like SSA4_0 highlights the potential of toehold switches as precise, customizable tools for controlling gene expression in response to endogenous cellular states, which marks another promising step forward for synthetic biology applications in therapeutic and research contexts.

Regarding the third sub-module of the trigger segment selection module, in which we use the trigger segment currently tested against its complement sequence, ideally it should be tested against the whole switch (with at least the 5’ UTR switch prefix and the early codons of the expression gene), however, in that stage the switch is not generated yet. This manifests a cyclic dependency pattern of the switch generation on the trigger segment selection and vice versa. Such dependency pattern could be solved by performing a few cycles until the convergence of a certain defined global generation metric score, thus implementing an end-to-end optimization process. This could only be solved in the future, if enough data is collected which will enable learning score function and parameters for trigger segment selection and toehold predicted efficiency score.

An interesting question which should be raised is why toehold switches work in eukaryotes. We have not found a previous biophysical analysis trying to explain the experimental observation of achieving higher translation in the presence of a trigger, which is not trivial since in both off and on states a formed duplex should be unzipped. While in the off state the hairpin should be unzipped for translation initiation, in the on state the duplex formed by the trigger and the trigger binding site should be unzipped as well, not to mention that in on state the duplex segment formed is longer comprising both the stem length and trigger binding site length. We have raised a few possible hypotheses which could explain the higher translation rates observed for the toehold on state in eukaryotes. Though the binding segment is longer for the intermolecular trigger switch interaction, the binding is destabilized by both structural constraints of the switch and the relatively long mRNA molecule of the trigger and by other molecules in the cell. The more influential part of the binding energy of the duplex is related to the stacking energy which achieves optimality when forming a certain helix structure which is usually interspaced when formed between different RNA molecules [35]. Another issue is the fact that in the presence of a trigger there is a competition between self-refolding into hairpin and the trigger switch branch migration reaction ending in the formation of trigger switch duplex. The described competing states can be assumed to lower the effective binding energy (Fig. 5A).

The ability to design synthetic translational regulated switches for eukaryotes may remove the barrier for the development of many biotechnological applications and holds a promising therapeutic potential. Our tool exhibits a great improvement toward the design of more efficient toehold switches for eukaryotes yet holds a potential to be tested in prokaryotes as well. The new architecture we presented suggests that diverting from the “canonical” design can reveal superior switch architectures. We believe that despite the much higher complexity of the molecular biology processes in eukaryotes obscuring a direct solution for achieving high dynamic range for translation regulated toehold switches in eukaryotes, further application of novel RNA switch design architectures, learning from high throughput data results and implementing the new understandings via enhanced computational models will lead to even higher dynamic ranges when activated by any endogenous trans RNA sequences.

## Methods

### Biological Material and methods

YEAST: YPD (yeast rich medium) - 1% Bacto yeast extract, 2% Bacto peptone, 2% Glucose. For solid media 20 g/L agar was added.

SD (yeast defined medium) - 0.67% Bacto yeast nitrogen base w/o amino acids, 2% Glucose. Amino acids were added according to requirement. For solid media 20 g/L agar was added.

SM (yeast sporulation media) - K-Acetate- 20g

BACTERIA: LB - 2% Bacto LB extract. Ampicillin (Amersham) 50 mg/L was added to LB+Amp plates. For solid media 20 g/L agar was added.

**Yeast strains:** Unless differently stated, strain used in the experiments is: BY4741: *MATa ura3Δ0 his3Δ1 leu2Δ0 met15Δ0*

Standard Yeast Molecular genetics techniques were used to delete individual genes.

**Plasmids:** unless stated differently, centromeric plasmid used were high copy number plasmids, 2µ pRS426 (*URA3*) or pRS425 (*LEU2*).

**Cloning:** standard cloning techniques were used, cloning *in vivo* on plasmids was done by Gap repair [36]. Toeholds were constructed using Twist Gene blocks assembled using Gap repair into pRS425 while the reported was expressed on pRS426 in the exogenous gene experiment. in the endogenous experiment the toehold with mCherry as expressed gene was assembled in to the pRS425 plasmid.

**Plate reader:**

<u>Exogenous experiment</u> - Cells were grown ON in 15ML tubes with selective media. On the morning of experiment, cultures were diluted 1:25 and grown until reaching an OD of 0.7-1 as measured by a spectrophotometer OD600. From this culture, 50 microliter was taken to an Eppendorf tube, spined down and then resuspended in 200 microliter PBS. Cultures were divided into separate wells in a 96 wells plate and then immediately measured for fluorescence and OD600 again. Fluorescence was normalized to the OD as measured by plate reader and controls of a WT strain and wells with only PBS were measured at every different plate to ensure the robustness of the results.

<u>Endogenous experiment</u> - All Cells were grown 2X overnight on selective plates (SD-,URA-) at 25°C (Non-Induced State). Then plates were divided into two groups (each group containing all variants), the first continued to be grown for 2 more nights at 25°C, and the second was grown at 37°C. Following overnight incubations, A sterile pipette tip was used to collect a small portion of each colony, which was then suspended in 500 µL of PBS. Then cells were washed 3 times and resuspended in enough PBS to have 200 µL of solution for each plate reader well. Cultures were divided into separate wells in a 96 well plate and then immediately measured for fluorescence and OD600 again. Fluorescence was normalized to the OD as measured by plate reader. A few wells with a control of const. mCherry grown at 25°C were placed on every plate for later usage when all measurements within a plate are normalized to those measurements.

### Pipeline for computational design of toehold

There is an inherent trade-off between usage simplicity and generability, we implement some basic off-the-shelf architectures and already pre-adjusted parameters while allowing the flexibility in an advanced options section to tune broad range of parameters and customize the activation of an assortment of features.

We list the main features of our tool and elaborate the biological motivation and role in toehold design by our tool.

#### Basic toehold architecture for prokaryotes

Our tool enables design of toehold switch for both prokaryotes and eukaryotes. For prokaryotes we use the “canonical” architecture previously demonstrated to yield high on/off ratios [11], [12], which is a basic hairpin sequestering the RBS in the hairpin’s loop and the starting codon in a bulge at the hairpin’s stem (Fig. 1A).

#### Hairpin’s general measurements

Though, most of the research regarding toehold switch following the first demonstrated high on/off ratios in prokaryotes [11], [12] used the specific measurements of the hairpin’s parts which achieved the highest on/off ratio, we allow the user to tune them, including stem size, loop size, toehold binding site size and relative RBS or Kozak offset to the end of hairpin’s loop (Fig. 1A,B,C).

#### Remain closed stem

One of the reasons the “canonical” architecture for toehold switches for prokaryotes has a bulge against the start codon is the intent to avoid introducing a constraint on the trigger sequence which by design should be a reverse complement to the downstream stem with the start codon included. Since the RBS is located at the end of the loop and should be at least 5-6 nucleotides upstream the start codon this leads to a toehold switch with a stem no designed to be fully opened by the trigger.

In our architectures for eukaryotes, we have no such constraint when placing the Kozak sequence after the stem and for the architecture implementing Kozak in loop, there is no spacing between the Kozak sequence and the start codon (the start codon is part of the Kozak sequence), so there is no inherent design resulting in few non opened stem nucleotides in activation. Nevertheless, we have added such an option to add few unopened nucleotides for eukaryote toehold switch design, according to the research that shows that mild or weak secondary structure binding near the start codon can enhance translation initiation rate [37]

#### Novel Kozak location architecture for eukaryotes

Previous research regarding toehold switches for regulation of canonical translation in eukaryotes applied the same architectural method used successfully for prokaryotes involving the sequestration of the initiation critical elements in the toehold’s stem-loop, just replacing the RBS with Kozak sequence [13]. As mentioned above, these attempts resulted in a poor on/off ratio.

One of the novelties of our work is the presentation of new toehold switch design architectures demonstrating a method change regarding the placement of the Kozak sequence. We added a new architecture placing the Kozak sequence after the designed hairpin structure followed by the expression gene (Fig. 1C). In such a design strategy, we also avoid the need for adding a linker.

For eukaryotes the user can choose between the architecture placing the Kozak either in the hairpin’s loop (Fig. 1B) or the new one placing it after the hairpin’ stem (Fig. 1C).

#### Novel Kozak post stem distance feature for eukaryotes

The novel positioning of the Kozak sequence after the hairpin’s stem has led us to another novel feature of placing the Kozak sequence at some distance after the stem (Fig. 1C) enabling optimization of the pre-Kozak sequence in such a way that moves the Kozak and the expression gene further downstream creating an unconstraint gap and decreases the local binding affinity of the trigger binding site (expected optimally to be a single-stranded segment ready for the trigger) to the Kozak sequence and expression gene (Fig. 5A).

#### Novel Pseudoknots avoidance

Though some RNA secondary structure prediction libraries include pseudoknots prediction, no such support is available for the selected inverse problem solver to allow the design of a mRNA sequence while declining secondary structure candidates containing significant pseudoknots. We have added an algorithmic layer that allows us to prefer a significant pseudoknots-free structures opening the possibility of testing another novel feature, whether avoiding pseudoknots results in a more efficient toehold switch design (Fig. 5B).

#### Novel Dual expression gene codon bias

Another novel feature related to toehold design is the codon bias optimization of the expression gene. Codon bias optimization of expression gene codon is optionally performed in two different stages. It is first optionally performed at the pre-toehold generating stage in which all the expression gene’s codons are optimized for high translation rate according to CAI (Codon Adaptation Index) score algorithm [25] using CAI reference scores pre calculated for the specified organism (Fig. 5C). Codon bias is also optionally used later in a different manner in the toehold switch generation phase, in which alternative codons for each of the early expression gene’s codons are allowed to be optimized in the switch design phase as an added degree of freedom to achieve a higher dimension of search space of designs resulting in a higher predicted metric score (Fig. 5A). In this dual stage codon bias optimization, we aim to achieve both higher translation rates as well as higher predicted metric score. Adding more degrees of freedom to search space may allow achieving more resemblance of the designed switch structure to that of the desired target structure, which may have even more importance in the segments which should be single-stranded.

#### Novel Kozak guided early reporter gene positioned based codon optimization

The mentioned translation optimization process applied on the reporter gene using CAI based algorithm does not consider the codon position nor its neighboring codons context, however, Kozak sequence and nearby position PSSM (position-specific scoring matrix) [38] statistics and experiments exhibit a tremendous influence on the translation efficiency. We implemented another layer of position-based codon optimization for the early ORF (open reading frame) codons in proximity to the start codon of the reporter gene to favor high score codons in accordance with a pre-built per organism PSSM table of those neighboring codons. The statistics are generated using highly expressed genes data.

#### Linker and Trigger mRNA codon bias

Our tool enables codon bias high translation optimization for the linker sequence in those architectures it is used and for the trigger mRNA sequence in cases where non endogenous mRNA is used, and user has biological experiment control on the production of the trigger mRNA involved.

#### Optimized triggering mRNA segments selection

User may either specify a pre-selected trigger or apply another preliminary trigger calculation stage, in which a trigger mRNA is specified, and our tool runs an optimization algorithm for selecting the best trigger segments to be used for that specified mRNA molecule (Fig. 1E).

Calculation is performed by our algorithm for finding low expected probability of self-folding segment (see more details in the sub-section Algorithms). We also allow the selection of multiple optimal trigger segments candidates and enable filtering the search space by setting required GC ratio range.

#### Conserved sequences customizations

Our tool lets the user use pre-defined conserved sequences optimized for different organisms, including *E. coli* and *B. subtilis* for prokaryotes and *S. cerevisiae*, *K. pastoris*, *CHO* and *HEK293* cell lines for eukaryotes. The conserved sequences are the RBS or Kozak and the Linker. We also enable a customization mode in which the user can provide those conserved sequences for other organisms not supported yet in our tool.

#### Repeated nucleotides avoidance

Repeated RNA nucleotides segments have in some cases molecular biology meaning, for example guanine tetrad and when few of them are stacked on top of each other forming a G-quadruplex are motifs with certain tertiary structure and many biological roles. Those segments have also a manufacturer synthesis limitation when ordering RNA sequence with high level of such repeated segments. We allow the user to reject switches with nucleotides repentance above a set threshold.

#### Off-target detection

Our tool provides a feature for the detection of similar endogenous RNA sequences in the genome of the selected organism to each of the trigger segments candidates selected in the second module. The tool returns the RNA molecule with the highest matching similarity combined from sub fragments compared against the tested trigger segments. The algorithm returns a detected potential off-target only if its identity is above a threshold which is assumed to be high enough to initiate the branch migration reaction with the toehold switch’s trigger binding site and resulting in full hybridization and undesirably activation of the switch. This is a simplified implementation neglecting the abundance of the similar RNA molecule which should be more relevant to its actual influence and not testing if it has a similar differential expression pattern to the original trigger mRNA in which case it will not be considered an off target. The low-level calculations are performed using the BLAST package [39].

#### In vivo immune response uridine depletion and epitope resemblance report

Our tool advanced options section offers some features regarding the generation of toehold switches intended for testing their in-vivo usage potential. The tool includes an implementation of uridine depletion codon bias adaptation algorithm to lower immune response [40] and it also enables running a search algorithm for epitopes against an appropriate epitope database (www.iedb.org).

#### Visual report

Our tool returns the results of multiple optimized generated toehold switch candidates for each trigger segment selected in a CSV tabular file. These fields include generation parameters and some metadata fields. It includes calculated model related detailed scores for structural and concentration predicted optimality, estimated switch-switch homo dimer binding energy, off-target description if detected in case specified to check by user (an elaborated explanation is available in the tool’s site user guide page).

Another CSV file is attached with all ORF sequences including the source and optimized versions of the expression gene as well as the trigger gene and linker if used.

Finally, for the candidate with the highest predicted metric score [41], according to the core algorithm used for designing nucleic acids [42], among each group of candidates for a selected trigger, a thorough visual report in pdf format is generated.

The visual report includes a summary of all generation parameters, metadata and detailed calculated model related scores from the CSV file. It also contains two dot plots. One dot plot is the toehold switch self-folding base pair probabilities matrix. An example is shown in Fig. 6A, in which the blue square marks the switch hairpin’s stem segment. For the ideal design when the switch is without the trigger it should be in an off state achieved by the stem being base paired with high probability. As shown in the example the color of the diagonal line in the blue square is white which marks high probability of base pairing. There is some “leakage”, marked by lines outside the square with colors of less probability showing unwanted base pairing. The second dot plot is the toehold switch with the trigger co-fold base pair probabilities matrix. An example is shown in Fig. 6B, in which the red square marks the toehold switch binding site. For the ideal design when the switch is with the trigger it should be in an on state achieved by the trigger binding to the binding site and opening the stem. As shown in the example the color of the diagonal line in the red square is white which marks high probability of base pairing of the trigger with the binding site. The stem should be opened except for the remain stem size (explained above), and as can be seen the stem segment in blue square has no diagonal line since the stem is predicted to be successfully opened except a small line at the edge marking the stem remain size segment.

**Fig. 6:**
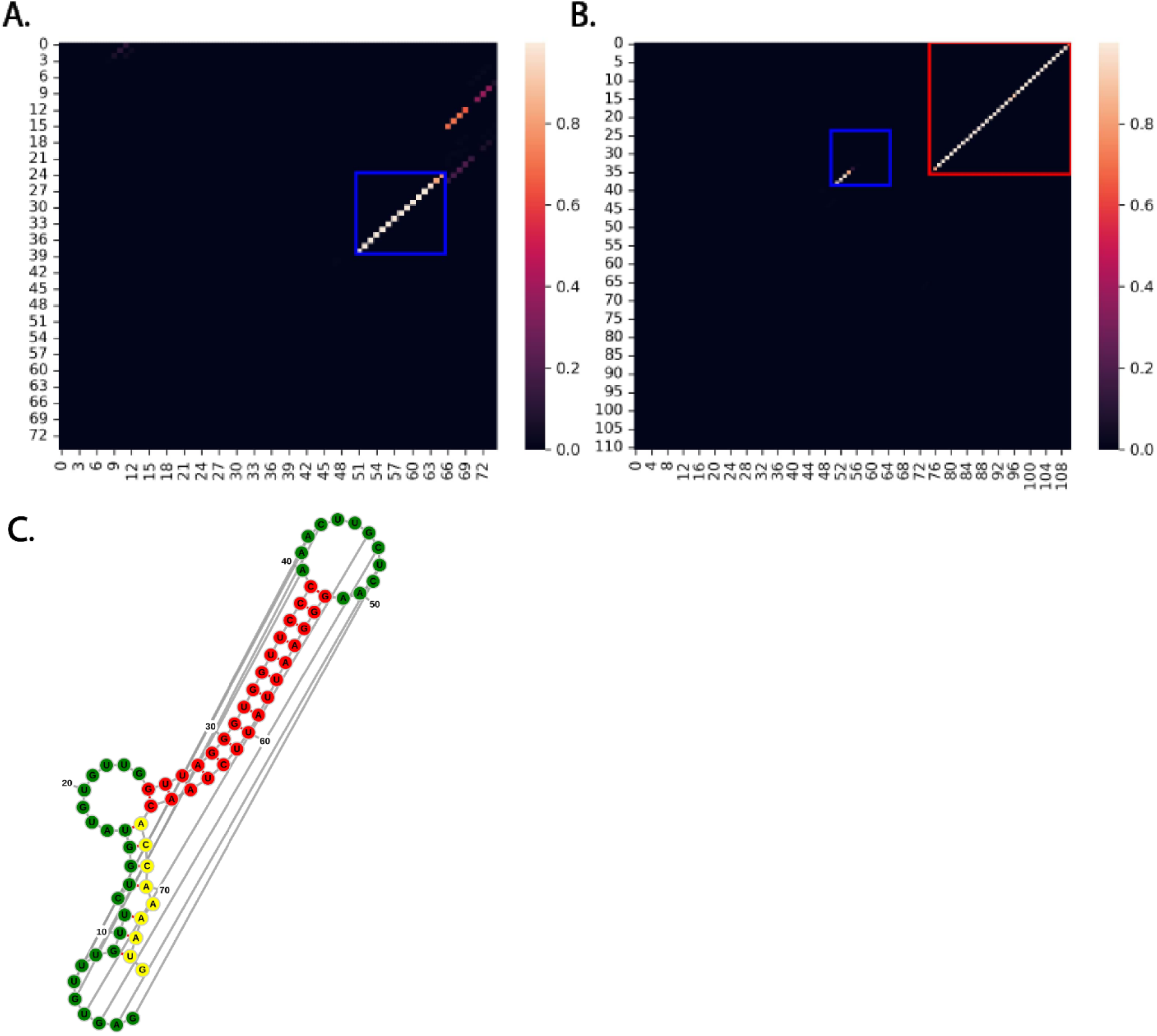
Visual report. **A.** Toehold switch co-fold base pair probabilities dot plot. The blue square marks the toehold hairpin’s stem segment. **B.** Toehold switch with trigger co-fold base pair probabilities dot plot. The red square marks the toehold’s binding site segment. **C.** Switch MFE structure with pseudoknots.

The visual report contains also an illustration of the MFE secondary structure including pseudoknots of the switch RNA sequence (Fig. 6C) using the Forna tool [43].

### Algorithms

We Divide the toehold switch design engineering problem into 3 major modules (Fig. 1E).

#### Trigger mRNA selection module

The first module is the selection of the trigger mRNA molecule. In the perspective of an appropriate triggering RNA molecule for the activation of a bi-stable on-off toehold switch optimized for differentiation of a certain organ, a certain cell type or cell state, the ideal selected candidate should be the most differential endogenous gene expression compared between the two sets of cells, one in which the switch should be activated and the other in which the switch should remain in off-state. Those differentially expressed genes should be the best candidates for gaining the highest on-off ratio and. Among those differential genes another criteria should be considered for preferring those genes with a high absolute expression in the on state. To achieve that we need an RNA sequencing database of the target organ or cell type for the two differential states. Since in general the application could vary for different users’ needs and task definition, each requiring access to different database types, not to mention their dynamic evolvement, we did not integrate such capability. Users may use any available tool (e.g. DESeq2, edgeR) upon any data which fits its application. In the future we may implement such module into our pipeline, but for now our tool assumes the user provides it with either pre-selected trigger mRNA sequence for the further selection of optimal trigger segment locations within it or an already pre-selected such trigger RNA segment.

#### Trigger segment selection module

The second module is the module for selecting proper sub-segments within a pre-selected mRNA trigger molecule to serve as the trans-binding RNA to activate the toehold switch upon binding.

The assumption is that the most influential part on the capability of an mRNA segment to bind to another complementary RNA segment (the binding site in the switch is designed to have base pair complementarity to the selected trigger sub segment) is its structural accessibility for hybridization interaction to occur when the two molecules reach proximity. Under the simplification of neglecting other molecules interactions including small ligands, RNA binding proteins, and specifically other RNA-RNA interactions, the availability for physical RNA-RNA interaction is governed by self-folding tertiary structure of each of the interacting molecules before the interaction. Since the RNA molecule secondary structure provides a good approximation for physical availability while being algorithmically and computationally well explored, our algorithm relies on self-folding ensemble statistics of the secondary structure.

Thus, our model uses secondary structure prediction models to calculate an estimation of self-binding mRNA with itself to find the least self-bonded segment of a specified length within the trigger mRNA molecule.

mRNA molecules have a typical length of a few hundred nucleotides, however, the mRNA structure prediction algorithms based on free energy have exponential accuracy degradation with size. Moreover, the nature of mRNA molecule dynamics is being translated by ribosomes during its lifetime, so the reasonable hypothesis is that the mRNA never reaches global equilibrium and was demonstrated to have less stable long range secondary structure [44] not to mention that even without ribosomes the secondary and tertiary structure of RNA in general are unstable. According to these assumptions, our algorithm aiming to calculate an expected ensemble of locality-based approximation measurement is more appropriate.

Our trigger segment selection algorithm is combining scores of three sub modules’ scores. The first one integrates the data of single nucleotide folding probabilities in the segment, the second one integrates short segments folding probabilities within the candidate segment, and the third checks the thermodynamically favorability of such interaction to occur.

The naïve approach would be to perform a direct calculation of the whole mRNA free energy and self-nucleotides binding probability matrix, however as explained above a more appropriate approach we use is to perform many such calculation on a moving sub window and finally return a weighted average of the probabilities of each sub window.

For the first sub module, integrating single nucleotide folding probabilities, we integrate data from multiple sub windows (algorithm’s outer window) single nucleotide probabilities in which the candidate segment (algorithm’s inner window) is contained, using certain padding to avoid sub windows containing the trigger segment near the edge and moving the sub window with certain step size (Fig. 1D). Finally, we average them with a weighted sum giving weight proportional to the trigger segment’s proximity to the sub window’s center. The main parameter of the algorithm is its outer moving window size. Its size has a molecular biological meaning of segment which can be considered to achieve relatively local equilibrium. This segment size could theoretically be optimized if enough experimental data was present. We set a default value of around 120 which is a few ribosome footprints. The low-level calculations using ViennaRNA package [7] are performed once for each such moving window but used for different trigger segments contained within it.

For the second sub module, integrating short sub segments folding probabilities, we select an outer window size (same as the one used for the first sub module) for which inner window of the trigger segment is contained at its center. The algorithm further divides the candidate trigger segment into shorter sub segments of a constant size, we call seeds, with some overlap. For each candidate trigger segment, we average the probabilities of those short seed segments contained within the trigger segment. The seed size symbolizes physically the shortest segment we assume is needed for a hybridization base pairing interaction to initiate in a similar way it is used in some other packages (e.g. IntaRNA). The low-level calculations are performed using the Raccess library [45].

The third sub module calculates for each trigger segment the thermodynamical change of free energy between the interacting complex of an outer window containing in its center the tested candidate trigger segment with the trigger segment’s complement sequence against the sum of each of those components self-folding energies. The low-level calculations are performed using the IntaRNA package [35], [46].

#### Toehold generation module

The third module is the core module introducing the ability to design a toehold RNA switch molecule that has B-stable states, default off state, and an on state in the presence of a target trans-RNA segment provided to it as an input from the second module.

To design RNA molecules, with sequences resulting in a desired target structure we need to solve the inverse problem of RNA structure prediction. We have used the NUPACK library [42] as a core RNA design inverse problem solver. Though it has some built-in mechanisms for adding constraints some of our new features had to be implemented as another layer.

The core process algorithm is composed of some other sub-processes (Fig. 1F). As a preliminary stage we first optionally codon optimize the expression gene and linker if used and the early codons following the start codon in case of Kozak in eukaryotes.

We then proceed with generation of a few candidates of toehold switches using different random seed values due to the intrinsic randomness of the generation core algorithm.

Regarding the core toehold generation process there are few special methods which we elaborate on, but we first define the toehold generation metric score.

#### Ensemble Defect

According to thermodynamics and statistical mechanics theories, a complex of molecules in equilibrium does not reach a single state, but forms a dynamic system having many possible states, each with a different energy and distributed by the Boltzmann distribution which are defined as the ensemble of states. The core packages used for RNA structural prediction are based on dynamic programming algorithms using free energy models which return structural predictions results in term of probabilities. For self-folding RNA or RNA-RNA interactions we calculate the probability matrix of the nucleotide-nucleotide base pair probabilities. When modeling the interaction of some molecules, the resulting concentration of each of the reactants and products in the reaction can also be predicted based on each sub-complex energy.

For the inverse solver, we define a target structure. For an optimization process to run we must define a metric score. For the structural problem, some kind of distance function score is defined, defaulting usually to the normalized difference between sum of nucleotides which are correctly paired according to the target structure and those which are not. This score measuring the deviation of a structure from the ideal target is termed the defect score. The expectation of this score can be calculated for the entire ensemble of possible structures, weighted according to the Boltzmann distribution of each (set appropriately to each structure’s free energy). This whole ensemble score is termed as the structural ensemble defect [41]. The ensemble defect can also be calculated for the change of reactants and products concentrations from a certain defined target concentration level, which is termed concentration ensemble defect. The core inverse solver package we use aims to optimize the total ensemble defect which is the sum of the structural and concentration ensemble defects.

#### Pseudoknots avoidance algorithm

Since the core used package for RNA inverse solver design does not account for pseudoknot structures, we added a layer which would seek to remove substantial pseudoknot. The algorithm checks the predicted structure with a forward solver that includes pseudoknots in its prediction and in case of the presence of substantial ones, dynamic constraints are added to the core inverse solver and the process is iterated until stopped by either success significant pseudoknots removal or some other stop criteria (Fig. 1F). In the case of our hairpin structure, pseudoknot removal aims to avoid RNA-RNA interaction like the loop-linear interaction between the pre-stem or post-stem segments and the loop (Fig. 5B).

Since the ensemble defect, which is the optimization metric used by core algorithm, is based on an ensemble of structures not including pseudoknots, the removal of such structures with significant pseudoknots will not only be invisible to the used metric score, but also may lead to higher defect scores since it narrows the theoretical solutions search space.

#### Lower Ensemble Defect Promotion method

For each set of parameters for the task of designing a toehold switch there is a certain reflected set of constraints and an appropriate set of degrees of freedom and a resulting search space. As mentioned, we use codon bias twice. Once in the preliminary stage on all ORF codons (expression gene’s ORF and trigger’s and linker’s ORFs if used) as codon bias optimization for higher translation throughput, and again in the core toehold design phase on few of the early codons of the expression gene as a constraint relaxation method not aiming for high translation throughput codons but for minimization of the ensemble defect achieved by allowing any synonymous set of codons replacement which will result in the lowest ensemble defect. This method should allow a direct improved avoidance of unwanted base pairing which are visible to the core engine (non-pseudoknot unwanted base pairing) like the linear-linear pre-stem and post stem interactions between the toehold binding site and the early codons of the expression gene (Fig. 5A). It has also indirect supporting influence on the pseudoknot avoidance algorithm through its repetitive iteration algorithm by introducing a wider search space.

#### Gene optimization

In the preliminary stage the trigger gene, the expression gene and the linker, if used, are optionally optimized for higher translation. Codon Adaptation Index [47] scores are pre-calculated for few organisms and the appropriate data according to the selected organism is used. Our algorithm does not implement naïve highest CAI score codon selection for each codon but adds randomness by selection of each codon according to a codon probabilities table which is built upon the CAI scores.

#### Uridine depletion

An advanced feature for in-vivo usage is optionally performed in the preliminary phase in which synonymous codons are used at the same optimization phase for optional higher translation optimization. If both codon bias optimization for higher translation and uridine depletion are specified, algorithm chooses the highest CAI codon by an appropriate organism’s codon probabilities table as mentioned above, however, after the algorithm’s first pass selection it will perform a reselection process for each codon which has a synonymous codon with a lower uridine count. If only uridine depletion is specified, the codon with lowest uridine is replaced in case it is not the already the set codon. This is performed on the expression gene and the linker in case it is used.

#### Epitope avoidance

This is more relevant when we add a linker before the gene of interest unless the expression protein is de-novo one. We return a flag whether the ORF including the stem after the start codon, the linker, and optimized expression gene have resulted in a sub-sequence which is classified as an epitope according to an epitope database (www.iedb.org).

#### Coupling between sub-modules

The three high-level modules of the pipeline (Fig. 1E) have an inherent coupling. As explained above we decided not to include an implementation of the first stage of choosing an mRNA, thus relying on users’ pre-selection of an appropriate mRNA according to their criteria and preferences. In that operation mode we assume that a user has selected for example a differential mRNA trigger that is highly expressed in the target cell type that the toehold switch should be activated while having low expression levels in other non-target cell types. However, optimizing the first stage by itself is not ideal since there exists a strong reciprocity with the second stage of selecting a segment within that already pre-selected mRNA. The span of the search space for optimized segment as unfolded sequence to serve as a toehold trigger is limited to that already pre-selected single mRNA. Moreover, strong coupling exists also between the second stage of trigger segment selection and the third stage of toehold generation, as for example the best selection of the best segment as trigger according to unfolded metric may be less optimal for toehold generation when having high reverse complementariness similarity with a segment within the expression gene. Theoretically the ideal strategy for implementing such a pipeline would be using an End-to-End optimization, however, we have chosen a simple design implementation of optimizing each stage separately since not enough prior data was available for training a model which could intelligently combine the different metric score of each stage and assign them relative weights appropriately. As already mentioned in the introduction there are a few previously developed computational tools for toehold generation, for example a tool [22] which is using a propriety prediction score based on a model trained on 181 toeholds for selection of the optimal one. The tool’s major drawbacks are that both the training and the generated toehold switches are limited to a specific architecture with a very limited degree of freedom in tuning parameters and trying new designs, not to mention being already biased for prokaryotes and not learning transferable when applied to eukaryotes designs.

### A user-friendly web tool

Our web tool user interface is implemented in a single form (Fig. 4A,B). Since different architectures require different parameters, we have implemented a dynamic form changed upon user selections. The first required parameter is the users email address for which results will be sent to upon background calculation process completion. Then the user selects a domain, either prokaryotes or eukaryotes and a desired model organism from a list of supported ones. Users may either provide a pre-selected trigger segment sequence as a trigger or set an entire mRNA molecule to choose a trigger from; in the second case, its size will be determined implicitly by the sum two parameters in the advanced section, the trigger binding site size and the stem size. Users may instruct the selection algorithm to filter only segments with GC ratio range. The last required parameter is the expression gene.

For more advanced customizations a button to show advanced options exposes many design parameters. If the domain selected is eukaryotes, then user may select which architecture to use either placing the Kozak in loop or after the stem, and selecting between two architectural trigger binding location options, whether binding occurs before or after the stem. In the case of the prokaryotic domain, default RBS and linker sequences are provided but may be set to customized sequences. In all architectures, general measurements are allowed for customized change including stem size to be opened by trigger, remain closed stem size, loop size, trigger binding site size, RBS or Kozak start position offset and linker sequence and start offset in case it is used when RBS or Kozak is in the hairpin loop. A 5’ UTR prefix is enabled to be set, temperature and a set of flags, including whether to optimize the expression gene for translation, whether to optimize the expression gene’s early codons for higher predicted metric scores, whether to apply the pseudoknot avoidance algorithm, whether to perform uridine depletion, whether to impose reading frame and stop codons check and whether to avoid highly repetitive nucleotides.

The returned email message consists of a csv file with the data of all the generated toehold candidates which share the same parameters and only differ due to the intrinsic randomness of the generation algorithm. Each such set is generated for each trigger segment candidate selected if an explicit mRNA is provided. The email also contains visual reports in a pdf format. It is generated for the highest predicted metric score candidate for each of the trigger segments selected and has both textual summary of results metadata and calculated detailed model related scores as well as toehold switch with (Fig. 6A) and without (Fig. 6B) trigger dot plots of nucleotide binding probabilities matrices and RNA folding secondary visual structure illustration (Fig. 6C).

## Data Availability

The tool can be accessed in http://www.cs.tau.ac.il/~tamirtul/Triggate.

## Acknowledgments

We would like to thank Dr. Daniel Dovrat, the IGEM TAU 2022 group, and the IGEM TAU 2024 group, for helpful comments.

## Supplementary information

Supplementary file with supplementary figure 1: supplementary1.docx.

Supplementary file heterologous mRNA targeting experimental data: supplementary2.xlsx.

Supplementary file endogenous mRNA targeting experimental data: supplementary3.xlsx.

