## Supplementary information for "Computational design of toehold switches in eukaryotes and prokaryotes for efficient post-transcriptional control"

### Supplementary figure 1

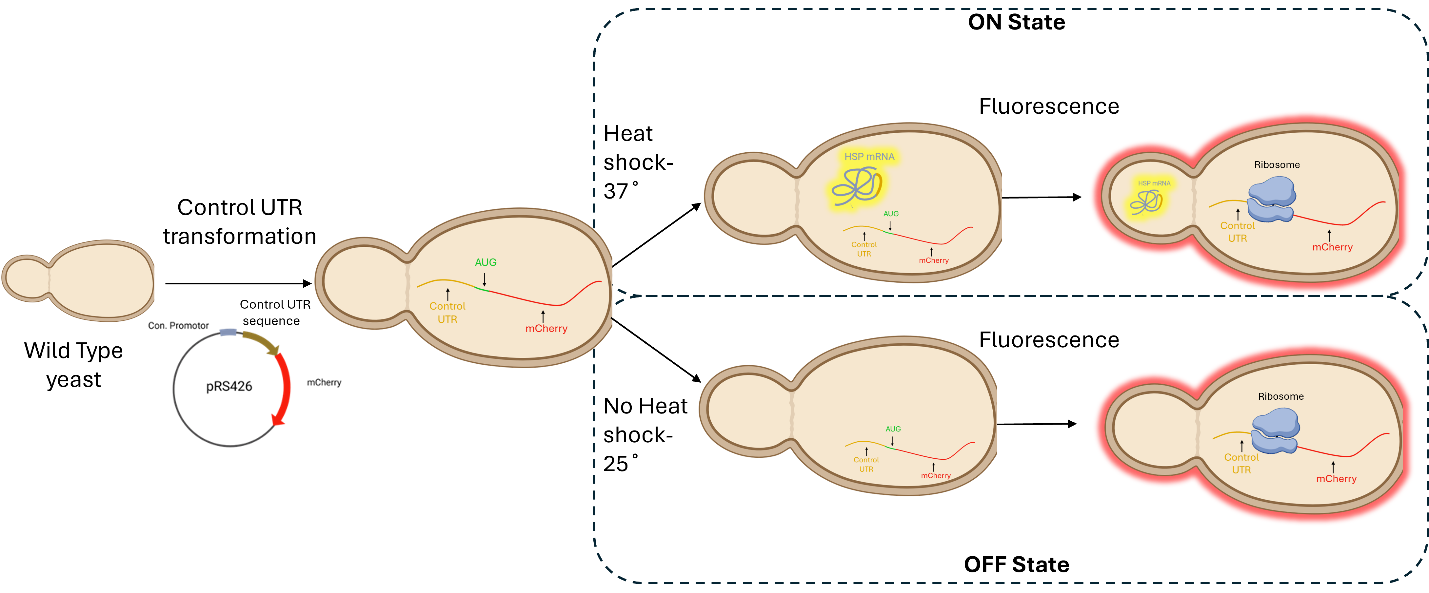

Supplementary Fig. 1**: Control UTR experimental design (results for this experiment are in figure 3C)**
